# Wikidata as a semantic framework for the Gene Wiki initiative

**DOI:** 10.1101/032144

**Authors:** Sebastian Burgstaller-Muehlbacher, Andra Waagmeester, Elvira Mitraka, Julia Turner, Tim Putman, Justin Leong, Paul Pavlidis, Lynn Schriml, Benjamin M. Good, Andrew I. Su

## Abstract

Open biological data is distributed over many resources making it challenging to integrate, to update and to disseminate quickly. Wikidata is a growing, open community database which can serve this purpose and also provides tight integration with Wikipedia.

In order to improve the state of biological data, facilitate data management and dissemination, we imported all human and mouse genes, and all human and mouse proteins into Wikidata. In total, 59,530 human genes and 73,130 mouse genes have been imported from NCBI and 27,662 human proteins and 16,728 mouse proteins have been imported from the Swissprot subset of UniProt. As Wikidata is open and can be edited by anybody, our corpus of imported data serves as the starting point for integration of further data by scientists, the Wikidata community and citizen scientists alike. The first use case for this data is to populate Wikipedia Gene Wiki infoboxes directly from Wikidata with the data integrated above. This enables immediate updates of the Gene Wiki infoboxes as soon as the data in Wikidata is modified. Although Gene Wiki pages are currently only on the English language version of Wikipedia, the multilingual nature of Wikidata allows for a usage of the data we imported in all 280 different language Wikipedias. Apart from the Gene Wiki infobox use case, a powerful SPARQL endpoint and up to date exporting functionality (e.g. JSON, XML) enable very convenient further use of the data by scientists.

In summary, we created a fully open and extensible data resource for human and mouse molecular biology and biochemistry data. This resource enriches all the Wikipedias with structured information and serves as a new linking hub for the biological semantic web.

## Introduction

Wikipedia (www.wikipedia.org) is a well established encyclopedia and collection of free form text, operated by the Wikimedia Foundation and edited by thousands of volunteer editors. As the seventh most-visited site on the Internet (http://www.alexa.com/topsites), Wikipedia has articles on a broad range of topics. With respect to molecular biology articles, at least two systematic efforts have been described, both initiated in 2007. The RNA Wikiproject created ~600 new Wikipedia articles on non-coding RNA families (1). In parallel, our Gene Wiki team created a collection of ~8,000 Wikipedia articles on human genes (2). Since its inception, the Gene Wiki has grown into an integral and strongly interlinked part of the English Wikipedia, now counting more than 11,000 articles (3, 4). The Gene Wiki articles have been expanded by the Wikipedia community and are highly accessed by users of Wikipedia, collectively viewed 4-5 million times per month.

Gene Wiki articles consist of two central parts, the free text representing a review of a gene’s function, biological role and impact on human health and disease, and the Gene Wiki infobox which provides structured data on the human gene and protein, and the orthologous mouse gene and protein. The data in the infobox comprises standardized identifiers and chromosome coordinates as well as functional annotation with Gene Ontology terms (5), structural information from the Protein Data Bank (PDB) (6) and tissue-specific gene expression (7) (Figure 3).

Wikipedia has proven a highly effective medium for collaboratively capturing unstructured text, but is technically lacking in facilities for authoring structured data. Several attempts have been made to better represent structured data within Wikipedia (8, 9). In late 2012, the Wikidata project (wikidata.org) was launched with the goal of creating an open, structured knowledge repository to complement and facilitate the unstructured content in Wikipedia (10). Like all other Wikimedia projects, Wikidata can be edited by anyone, and maintains an extensive version history for every item to allow for easy comparisons or reversions to past states. All content in Wikidata is licensed under CC0 (https://creativecommons.org/about/cc0) and therefore can be used by anyone without restrictions.

Wikidata consists of two entity types -- items (e.g. https://www.wikidata.org/wiki/Q7474 for Rosalind Franklin) and properties (e.g. https://www.wikidata.org/wiki/Property:P351 for NCBI Entrez gene ID) -- and every entity is assigned a unique identifier. Wikidata items and properties all have a label, a description and aliases. Every item record contains a list of claims in the form of triples. The subject of the triple is the Wikidata item on which the claim appears, the predicate is a Wikidata property, and the object is a date, a string, a quantity, a URL, or another Wikidata item. For example, the claim that Rosalind Franklin received her Ph.D. at the University of Cambridge is represented as Q7474 (Rosalind Franklin) - P69 (educated at) - Q35794 (University of Cambridge). Claims can be further amended with qualifiers (to indicate the context in which the triple is valid), and references can be added to indicate the provenance of the claim. The overall combination of a claim and references is referred to as a “statement”. A full description of the Wikidata data model can be found at https://www.mediawiki.org/wiki/Wikibase/DataModel/Primer.

A primary motivation for creating Wikidata was enabling easy accessibility by all language-specific Wikipedias. Now, statements about any Wikidata item can be displayed in the context of any Wikipedia item. This process is facilitated by “interwiki links” that establish connections between structured Wikidata items and the Wikipedia articles they are most closely related to. All major projects from the Wikimedia Foundation, including the language-specific Wikipedias, are linked to Wikidata using interwiki links. These links can be established between any existing Wikidata item and other MediaWiki content pages, e.g. Wikipedia articles.

Currently, a single Wikidata item can have an interwiki link to many different language-specific Wikipedia articles on the same topic. In return, this setup allows for all linked language-specific Wikipedia articles to access data from the central Wikidata item. Importantly, a Wikipedia article in a certain language can only have one interwiki link to a Wikidata item and a Wikidata item can only be linked to one single Wikipedia article in a certain language. This system has the advantage that data is stored once in Wikidata, and then made available for reuse across the entire Wikimedia universe.

Wikidata enables programmatic querying and access outside of the Wikipedia context. Specifically, Wikidata offers a Representational State Transfer (REST) API to easily perform structured data queries and retrieve Wikidata statements in structured formats. In addition, more complex queries are possible via a SPARQL Protocol and RDF Query Language (SPARQL) endpoint (https://query.wikidata.org) and a custom-built WikiData Query (WDQ) tool (http://wdq.wmflabs.org/wdq/).

In this work, we describe our efforts to migrate our Gene Wiki bot from English Wikipedia to Wikidata. This system offers significant advantages with respect to maintainability of the data, accessibility within the Wikipedia ecosystem, and programmatic integration with other resources.

## Database construction and usage

In collaboration with the Wikidata community, we decided to implement the representation of genes and proteins in Wikidata as separate Wikidata items. These gene and protein items are linked by the reciprocal properties ‘encodes’ (P688) and ‘encoded by’ (P702) carried by genes and proteins, respectively (Figure 2). Furthermore, orthologous genes between species are reciprocally linked by the property ortholog (P684) and also link out to NCBI HomoloGene (11) with the HomoloGene ID (P593). Homologous genes in the HomoloGene database and therefore also on Wikidata gene items share the same ID and can also be associated this way. The community discussion and decision process to establish this model, a very crucial mechanism in Wikidata, as well as Wikipedia, can be viewed here: https://www.wikidata.org/wiki/Wikidata_talk:WikiProiect_Molecular_biology.

**Figure 1:**
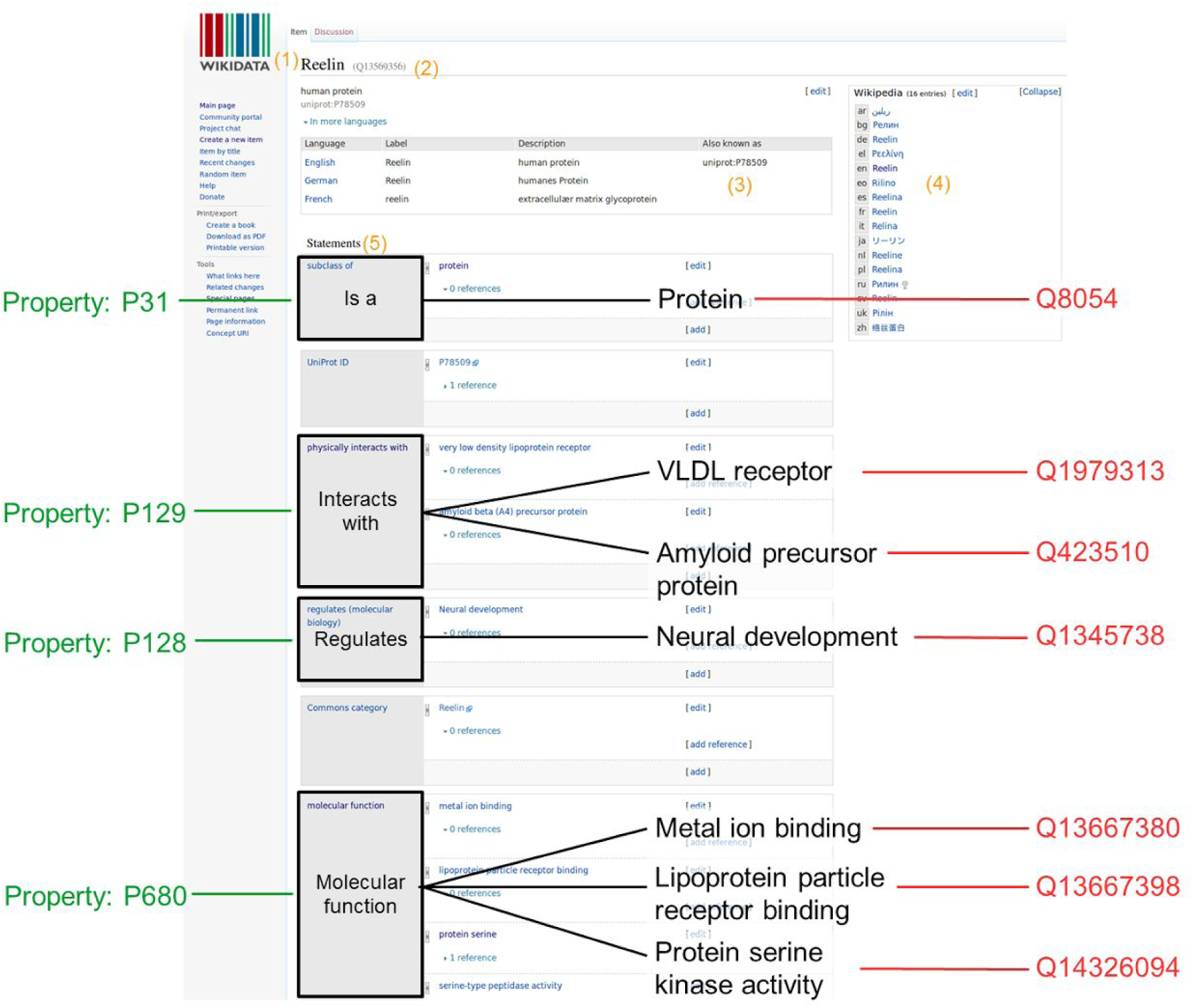
Wikidata item and data organization. Wikidata items can be added or edited by anyone manually. A Wikidata item consists of: (1) a language-specific label, (2) its unique identifier, (3) language specific aliases, (4) interwiki links to the different language Wikipedia articles or other Wikimedia projects, and (5) a list of statements.

**Figure 2:**
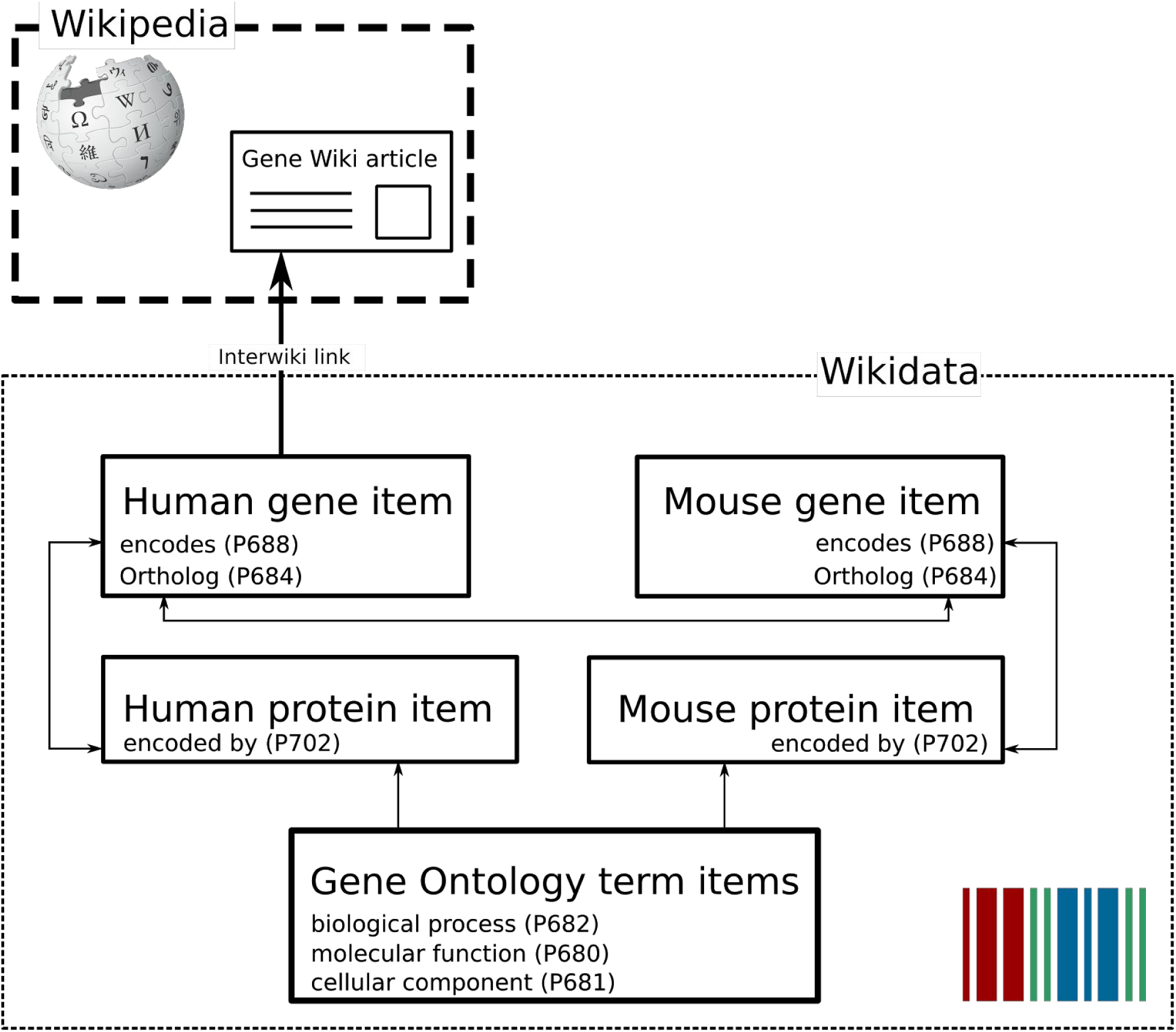
Gene Wiki data model in Wikidata. Each entity (human gene, human protein, mouse gene, mouse protein) is represented as a separate Wikidata item. Arrows represent direct links between Wikidata statements. The English language interwiki link on the human gene item points to the corresponding Gene Wiki article on Wikipedia.

**Figure 3:**
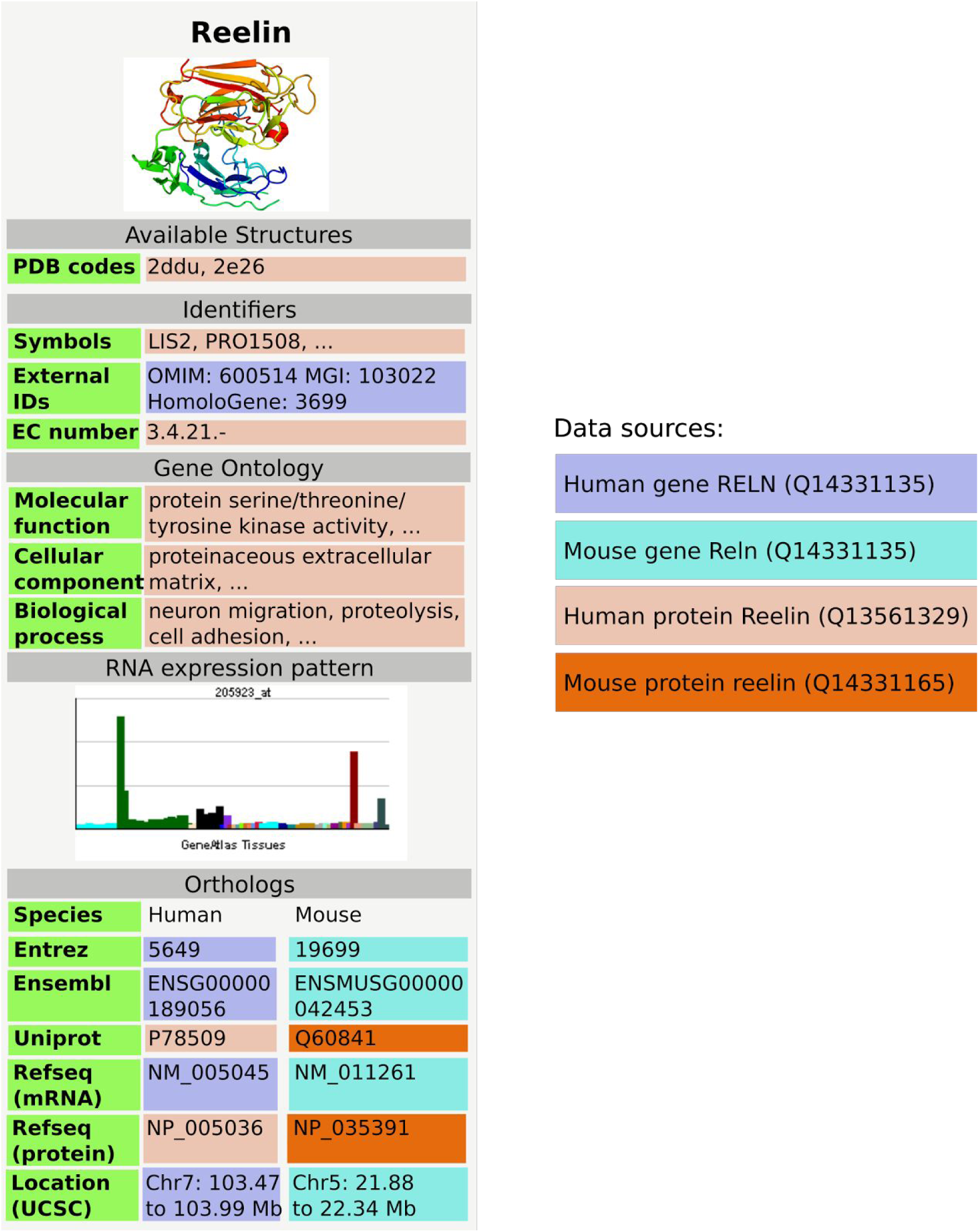
GeneWiki infobox populated with data from Wikidata, using data from Wikidata items Q414043 for the human gene, Q13561329 for human protein, Q14331135 for the mouse and Q14331165 for the mouse protein. Three dots indicate that there is more information in the real Gene Wiki infobox for Reelin.

### Data integration into Wikidata

We populated Wikidata with items for all Homo sapiens (human) genes, Homo sapiens proteins, Mus musculus (mouse) genes and Mus musculus proteins (Table 1). As described above, each gene and each protein was represented as a single Wikidata item. A full list of Wikidata properties used on gene and protein items is provided in Table 2.

**Table 1:**
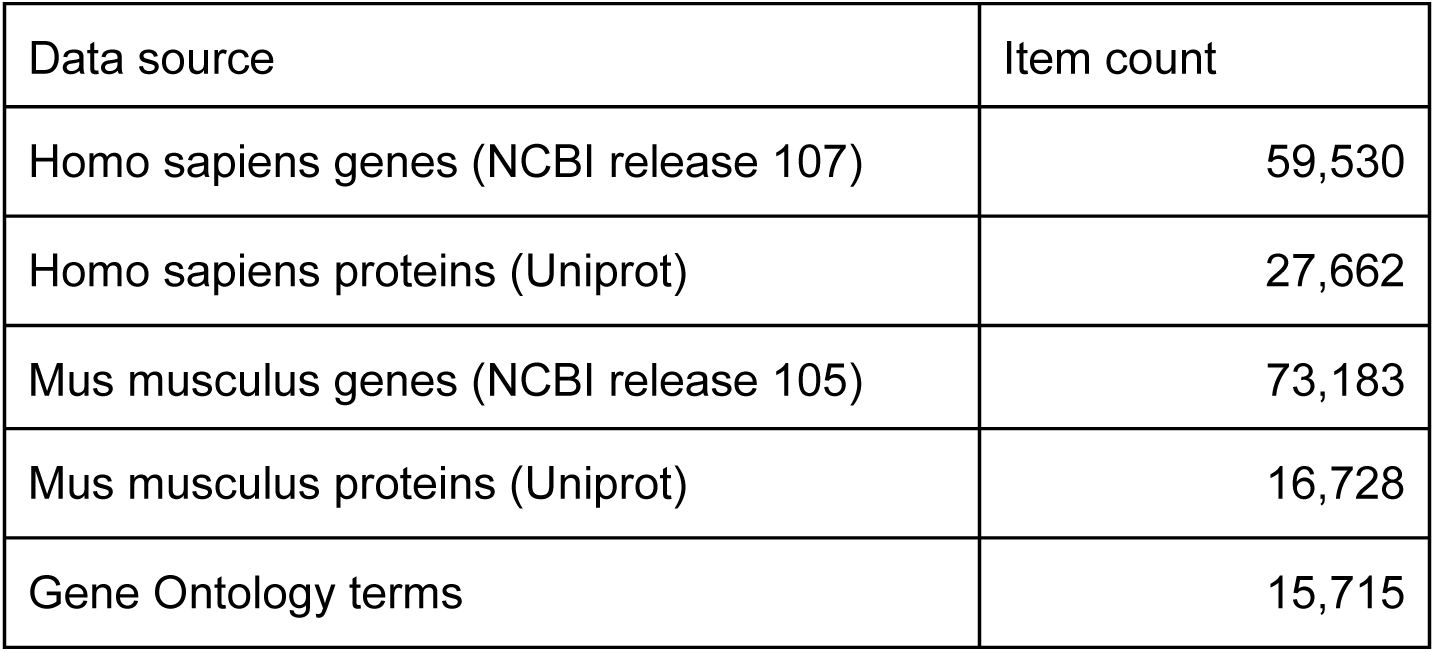
Overview on Homo sapiens and Mus musculus data in Wikidata.

**Table 2:** Wikidata properties used in this study. Column one contains the description as in Wikidata, column two the data type, column three the property number and column four a short description on the nature of the content.

| Label | Target data type | Property ID | Description |
| --- | --- | --- | --- |
| **Wikidata gene items:** |  |  |  |
| subclass of | Wikidata item | P279 | Defines to what category this item belongs to. Every gene item carries the value 'gene' (Q7187). Further subcategories are protein coding gene (Q20747295), ncRNA gene (Q27087), snRNA gene (Q284578), snoRNA gene (Q284416), rRNA gene (Q215980), tRNA gene (Q201448) and pseudogene (Q277338). |
| Entrez Gene ID | String | P351 | The NCBI gene ID as in annotation release 107 |
| found in taxon | Wikidata item | P703 | The taxon, either Homo sapiens (Q5) or Mus musculus (Q83310) |
| Ensembl Gene ID | String | P594 | Gene ID from the Ensembl database |
| Ensembl Transcript ID | String | P704 | Transcript IDs from the Ensembl database |
| Gene symbol | String | P353 | Human gene symbol according to HUGO Gene Nomenclature Committee |
| HGNC ID | String | P354 | HUGO Gene Nomenclature Committee ID |
| HomoloGene ID | String | P593 | Identifier for the Homologene database |
| NCBI RefSeq RNA ID | String | P639 |  |
| Chromosome | Wikidata item | P1057 | Chromosome a gene is residing on |
| Ortholog | Wikidata item | P684 | Ortholog based on the |
|  |  |  | Homologene database |
| Genomic start | String | P644 | Genomic start according to GRCh37 and GRCh38, sourced from NCBI |
| Genomic stop | String | P645 | Genomic stop according to GRCh37 and GRCh38, sourced from NCBI |
| Mouse Genome Informatics ID | String | P671 | Jackson lab mouse genome informatics database |
| encodes | Wikidata item | P688 | Protein item a gene encodes |
| **Wikidata protein items:** |  |  |  |
| subclass of | Wikidata item | P279 | protein (Q8054) |
| UniProt ID | String | P352 |  |
| PDB ID | String | P638 | Protein structureIDs from PDB.org |
| RefSeq Protein ID | String | P637 | NCBI RefSeq Protein ID |
| encoded by | Wikidata item | P702 | Gene item a protein is encoded by |
| Ensembl Protein ID | String | P705 |  |
| EC number | String | P591 | Enzyme Category number |
| Protein Structure Image | Wiki Commons Media File | P18 | Prefered protein structure image retrieved from PDB.org |
| Cell Component | Wikidata item | P681 | Gene ontology term items for cell components |
| Biological Process | Wikidata item | P682 | Gene ontology term items for biological processes |
| Molecular Function | Wikidata item | P680 | Gene ontology term items for molecular function |

Briefly summarized, for gene items, we imported data from the latest annotation releases from NCBI (Homo sapiens release 107, Mus musculus release 105) and created statements using many properties, including Entrez Gene IDs, RefSeq RNA IDs and chromosomal positions (11). Ensembl Gene IDs and Ensembl Transcript IDs were also imported and added to each gene item in Wikidata (12). Genes were categorized according to 8 subclasses. A generic subclass was used to identify a Wikidata item as a gene (Q7187), for increased granularity, the subclasses protein coding gene (Q20747295), ncRNA gene (Q27087), snRNA gene (Q284578), snoRNA gene (Q284416), rRNA gene (Q215980), tRNA gene (Q201448) and pseudo gene (Q277338) were added (Table 2). Genomic coordinates were encoded using the properties chromosome (P1057), genomic start (P644), and genomic end (P645), and the qualifier property ‘GenLoc assembly’ (P659) indicated the corresponding assembly version -- GRCh37 (Q21067546) or GRCh38 (Q20966585). Gene symbols were added based on the HUGO Gene Nomenclature Committee (HGNC) and HGNC IDs were also added to each gene item. For mouse gene nomenclature, the Jackson Laboratory Mouse Genome Informatics (MGI) data was used (13).

For protein items, we used UniProt as the the primary data source. All protein items received the ‘subclass of’ (P279) property value ‘protein’ (Q8054). A wide range of protein annotations were also added, including NCBI RefSeq Protein IDs (P637), Ensembl Protein IDs (P705), and PDB IDs (P638). Gene Ontology terms were added as Wikidata items, and annotations were added to protein items using three separate properties for Molecular Function (P680), Cell Component (P681) and Biological Process (P682).

We implemented this data importing process using Python (www.python.org) scripts, colloquially termed as bots by the Wikidata community. We run these bots with the Wikidata user account ProteinBoxBot (https://www.wikidata.org/wiki/User:ProteinBoxBot). The source code for the bots is available under GNU AGPLv3 (http://www.gnu.org/licenses/agpl.html) on our Bitbucket repository (https://bitbucket.org/sulab/wikidatabots/).

### Populating Gene Wiki infoboxes with data from Wikidata

As a first use case of the data, we focused on using the gene and protein data imported into Wikidata to populate Wikipedia Gene Wiki infoboxes. In our data model, we connected Wikipedia Gene Wiki pages to Wikidata human gene items with interwiki links (Figure 2). Four Wikidata items are required to fully represent one Gene Wiki infobox on a Wikipedia article. For this paper, we chose the gene *RELN* (protein Reelin) as an example. Specifically, the English language Wikipedia page (https://en.wikipedia.org/wiki/Reelin) of Reelin is directly linked to the human gene Wikidata item (Q414043) with an interwiki link. The Wikidata human gene item in turn links to the human protein (Q13561329), mouse gene (Q14331135), and mouse protein (Q14331165) using the data model described in Figure 2.

In order for Wikipedia pages to retrieve data from Wikidata, we used the MediaWiki extension module Scribunto (https://www.mediawiki.org/wiki/Extension:Scribunto), which integrates scripting capabilities based on the programming language Lua (http://www.lua.org). We created a new module in Lua code which generates the entire Wikipedia Gene Wiki infobox based on Wikidata data (https://en.wikipedia.org/wiki/Module:Infobox_gene). Using this new Wikidata-based infrastructure, a Gene Wiki infobox can be added to any Wikipedia page for a human gene by simply adding the markup code ‘{{infobox gene}}’, provided the Wikipedia page has a valid interwiki link to a Wikidata human gene item. This new system has been deployed on several test Gene Wiki pages within Wikipedia, and we will complete this migration after full community consensus is reached.

### Data usage beyond Wikipedia GeneWiki infoboxes

Data from Wikidata can be widely used in any application of interest. For example, the Gene Wiki infobox could now be rendered on any website on the Internet. As described in the introduction, a SPARQL endpoint, WDQ, the sophisticated Mediawiki API and a free text search engine constitute the main ways of querying data. These powerful facilities can easily be integrated in downstream analysis using Python, R or any other data analysis language which support seamless integration of data from the web. The SPARQL endpoint provides particularly powerful opportunities for dynamic data integration. For example, it would be useful to know: "Which genes, encoding for membrane proteins, are associated with colorectal cancer?". Content accessible through the Wikidata endpoint facilitates a single, distributed query that can answer this question immediately (Figure 4).

**Figure 4:**
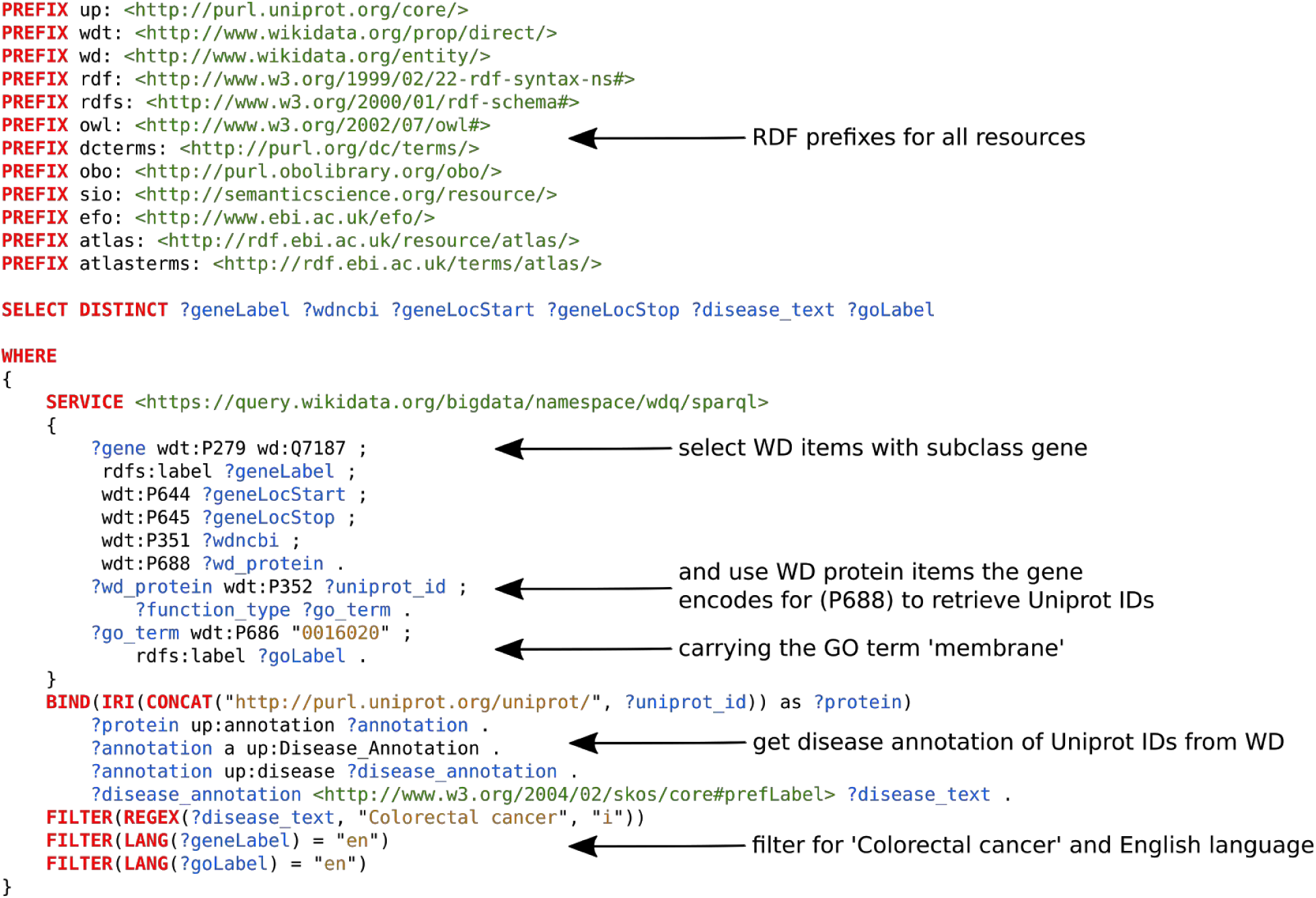
An example, federated SPARQL query, using the Wikidata and Uniprot endpoints. It retrieves all Wikidata (WD) items which are of subclass gene (Q7187), and encode for a protein (a separate WD item) which carries the Gene Ontology (GO) term ‘membrane’ (G0:0016020). The Uniprot ID of that protein is then used to complete the query on the Uniprot endpoint and retrieve the disease annotation and finally filter for the term ‘Colorectal cancer’. Colors: Red indicates SPARQL commands, blue represents variable names, green represents URIs and brown are strings. Arrows point to the source code the description applies to.

As the data is richly referenced, data origin and validity of statements can be reviewed instantly. Furthermore, the unique multilinguality of Wikidata, enabled by multilingual item labels, descriptions, and interwiki links to all of the Wikipedias allows the data to be used globally.

## Discussion

We created an open, community editable structured resource for all human and mouse genes and proteins using Wikidata as the technical platform. As a first use case of this data corpus, we demonstrated a remodelled Wikipedia Gene Wiki infobox which retrieves its data entirely from Wikidata, greatly simplifying the maintenance of these infoboxes. Until now, each of the 280 language-specific Wikipedias had to independently manage their infobox data in the context of MediaWiki templates meant for managing information display, not storage and retrieval, creating significant redundancy, a great risk for errors and inconsistencies, and out of date data. Now, not only do the Wikipedias benefit from higher data quality when based on a centralized data repository, Wikidata also benefits from the focused human effort in the global Wikipedia community.

In addition to these benefits to the Wikipedia community, the Gene Wiki effort in Wikidata also offers many data integration advantages to the biomedical research community. Biomedical knowledge is fragmented across many resources and databases, and small-scale data integration is an often-repeated exercise in almost every data analysis project. As for the language-specific Wikipedias, these small-scale integration efforts are incomplete, inefficient and error-prone. Instead, users now have the option of accessing and querying a central biomedical resource within Wikidata that is already pre-populated with many key resources and identifiers. While we certainly recognize that our effort does not yet include every resource in the biomedical space, Wikidata does empower any user to contribute data from their resource of interest. For example, any user could easily contribute data from other third party resources (e.g., International Union of Basic and Clinical Pharmacology (IUPHAR) (14), DECIPHER (15), COSMIC (16)) with minimal effort. These contributions can range from programmatic addition of large databases, to the output of medium-sized biocuration efforts, to individual statements added by individual users.

An important aspect of the broad applicability and reusability of Wikidata is its connection to the Semantic Web and Linked Open Data (Figure 4). Wikidata IDs give genes and proteins stable Uniform Resource Identifiers (URIs) in the Semantic Web, which in turn link to other common identifiers used in the biomedical research community. Moreover, Wikidata provides perhaps the simplest interface for anyone to edit the Semantic Web, which is otherwise limited by high technical barriers to contribute.

This work describes our initial effort to seed Wikidata with data from several key genomics resources. While this action has direct value to our Gene Wiki project, we hope and expect this first step to nucleate further growth of scientific data in Wikidata. With sufficient contribution and participation by the community, Wikidata can evolve into the most comprehensive, current, and collaborative knowledge base for biomedical research.

## Acknowledgements

We would like to thank the Wikipedia user RexxS for providing substantial help with the Gene Wiki infobox Lua code.

